# A single amino acid mutation in norovirus NS4 promotes viral spread

**DOI:** 10.1101/2025.08.27.672596

**Authors:** Mridula Annaswamy Srinivas, Robert C. Orchard

**Author notes:** **Correspondence to:** (R.C.O.).

## Abstract

Viruses can rapidly adapt and evolve to new, unfavorable environments due to their decreased replication fidelity, large reproductive index, and short life cycle. Often these adaptations that enable increased fitness in a new, specialized environment comes with a trade-off of decreased fitness in a standard, general environment. Understanding the tradeoffs of generalist and specialist viruses has provided important insight into vaccine development, mechanism of action of antivirals, and function of viral proteins. Here, we sought to identify how a specialist murine norovirus (MNV) could be converted to a generalist without a simple reversion of a genetic mutation. Previously, we found that a mutation in MNV (NS6^F182C^) overcame restriction by host protein Trim7 but decreased the efficiency of viral polyprotein NS6-7 cleavage and resulted in attenuation of this specialist virus. Here, we find that a single valine-to-isoleucine mutation in MNV non-structural protein NS4 (NS4^V11I^) is sufficient to rescue the attenuated replication of specialist NS6^F182C^ over multiple cycles of replication. However, NS4^V11I^ did not affect the defective polyprotein cleavage but instead the NS4^V11I^ mutation facilitates faster viral spread in vitro independent of interferon signaling. The emergence of this mutation in NS4^V11I^ suggests an unappreciated connection between NS4 and NS6 during norovirus replication and provides a system to define the unknown role of norovirus NS4 during infection.

**Importance:** Viruses and hosts are involved in a continuous arms race for survival. Often when viruses evolve to specialize in specific host environments, they lose their versatility and become specialists, only able to grow in one setting. This feature has been leveraged to create live-attenuated vaccines, identify the mechanism of action of antivirals, and to uncover fundamental aspects of viral replication. Here we aim to understand how a specialized virus can adapt again to become a generalist in the context of murine norovirus infection. We identify an unexpected connection between two murine norovirus non-structural proteins and uncover a role for the viral protein NS4 in viral spread. Taken together, these data provide new insight into viral evolution and the functions of norovirus proteins.

## Introduction

A fundamental question in virology is how viral fitness is tuned. As viruses evolve, they balance the selective pressures from host antagonism with constraints on the ability of viral proteins to perform their essential tasks. As viruses adapt to become more specialized in replicating in one environment, they frequently lose the ability to robustly replicate in other environments. In other words, adapted or “specialist” viruses tend to be less fit across multiple environments compared to their “generalist” precursors (1, 2). The molecular basis for the switch between specialist and generalist is largely unknown and challenging due to the large number of mutations accumulated in natural populations of viruses.

*In vitro* passaging of viruses enables a strong selective pressure to produce new specialist viruses. These systems have allowed for the discovery of antiviral targets, intrinsic barriers, inefficiencies to infection, and how viruses adapt to extreme environments (2). In many cases the fitness advantage of the specialist virus is limited to a specific set of conditions and the specialist virus is less fit than the generalist virus in alternative conditions. For example, selection of a ribavirin resistant Polio virus leads to a virus that can replicate significantly better in the presence of nucleoside analogs (3), but is significantly attenuated in mice compared to the wild-type virus (4, 5). Similarly, a Coxsackievirus B3 selected for increased replication speed was significantly attenuated in mice (6). Analogously, vesicular stomatitis virus (VSV) passaged onto HeLa or MDCK cells becomes less fit in the other host environment. However, alternating passaging enables equal fitness across hosts, demonstrating that not all adaptations come with trade-offs (7). Whether these specialist viruses can phenotypically switch to a generalist while maintaining the newly acquired mutation is not known. Such an investigation would provide information on the genetic interactions between viral proteins, likely giving molecular insight into evolutionary trade-offs.

Murine norovirus (MNV) is a leading model system to understand human norovirus infections in a natural environment. Human norovirus is the preeminent cause of gastroenteritis worldwide but is understudied due to the lack of a small animal model and limited replication of human norovirus in vitro. Noroviruses are non-enveloped, positive sense RNA viruses and are members of the Calicivirdae family. In a genome-wide screen for antiviral factors towards MNV, we determined that Trim7 expression induces a potent anti-MNV restriction phenotype (8). Trim7 binds and ubiquitinates the MNV non-structural protein NS6, which is the norovirus 3C-like protease (9, 10). Using an in vitro passaging approach, we determined that mutations that limit polyprotein processing of the NS6 and NS7 precursor (NS6-7) confer protection against Trim7 (9). Decreased processing of NS6-7 via introduction of a single point mutation (NS6^F182C^) provided a fitness advantage for MNV replication in Trim7 overexpressing cells (9). However, the replication of MNV NS6^F182C^ is severely attenuated in wild-type cells and in mouse models of disease (9). Despite the strong phenotype in vitro, Trim7 deficient mice had no differences in MNV replication or mounting an innate immune response to MNV (11).

Despite the lack of an in vivo phenotype, several lines of evidence suggest that the MNV-Trim7 system is an ideal experimental system to uncover fundamental aspects of norovirus biology and viral evolution. First, most viruses within the Caliciviridae family maintain their protease and polymerase as a fused precursor (the equivalent of NS6 and NS7 in noroviruses). It is unclear why these closely related viruses have different requirements for processing these proteins. Thus, investigating the consequences of a fused NS6-7 in MNV provides insight into Caliciviridae evolution. Second, our previously identified specialist virus MNV NS6^F182C^ provides a robust system to study fitness trade-offs in different environments (e.g. presence or absence of Trim7 overexpression). Third, advances in host and viral genetics enable facile genetic manipulation of both host and virus to study these questions in viral evolution.

Here, we used a forward genetic screen to identify a single suppressor mutation that enables MNV to replicate robustly even with inefficient cleavage of NS6-7. Unexpectedly, this mutation occurs in the viral non-structural protein 4 (NS4) consisting of a valine to isoleucine substitution. The NS4^V11I^ mutation not only improved the replication of a virus containing NS6^F182C^ mutation but also improved the replication of the wild-type MNV strain. NS4^V11I^ did not promote increased polyprotein processing but rather enhanced the spread of MNV independent of interferon signaling through STAT1. In summary, our data points to a genetic connection between processing of NS6-7 and MNV spread while also shedding light on the interconversion between generalist and specialist viruses.

## Results

### Viral evolution screen for attenuated CW3 NS6^F182C^

To try and convert the specialist virus NS6^F182C^ to become a generalist without losing the NS6^F182C^ mutation, we devised a passaging strategy in which NS6^F182C^ mutation on MNV strain CW3 (herein NS6^F182C^) was grown exclusively on Trim7 overexpressing BV2 cells (BV2-Trim7; **Figure 1A**). We initially passaged virus on BV2-Trim7 cells every 48 hours for 6 passages but observed minimal rescue. To increase the selective pressure, we limited the time of viral replication to 12 hours for passages 7 through 14. The passage 14 (P14) virus had a significant increase in viral replication over the original MNV^CW3^ NS6^F182C^ (P0) in wild-type BV2 cells (**Figure 1B**). Additionally, the replication of the P14 virus was similar to that of MNV^CW3^ wild-type (**Figure 1B**).

**Figure 1.**
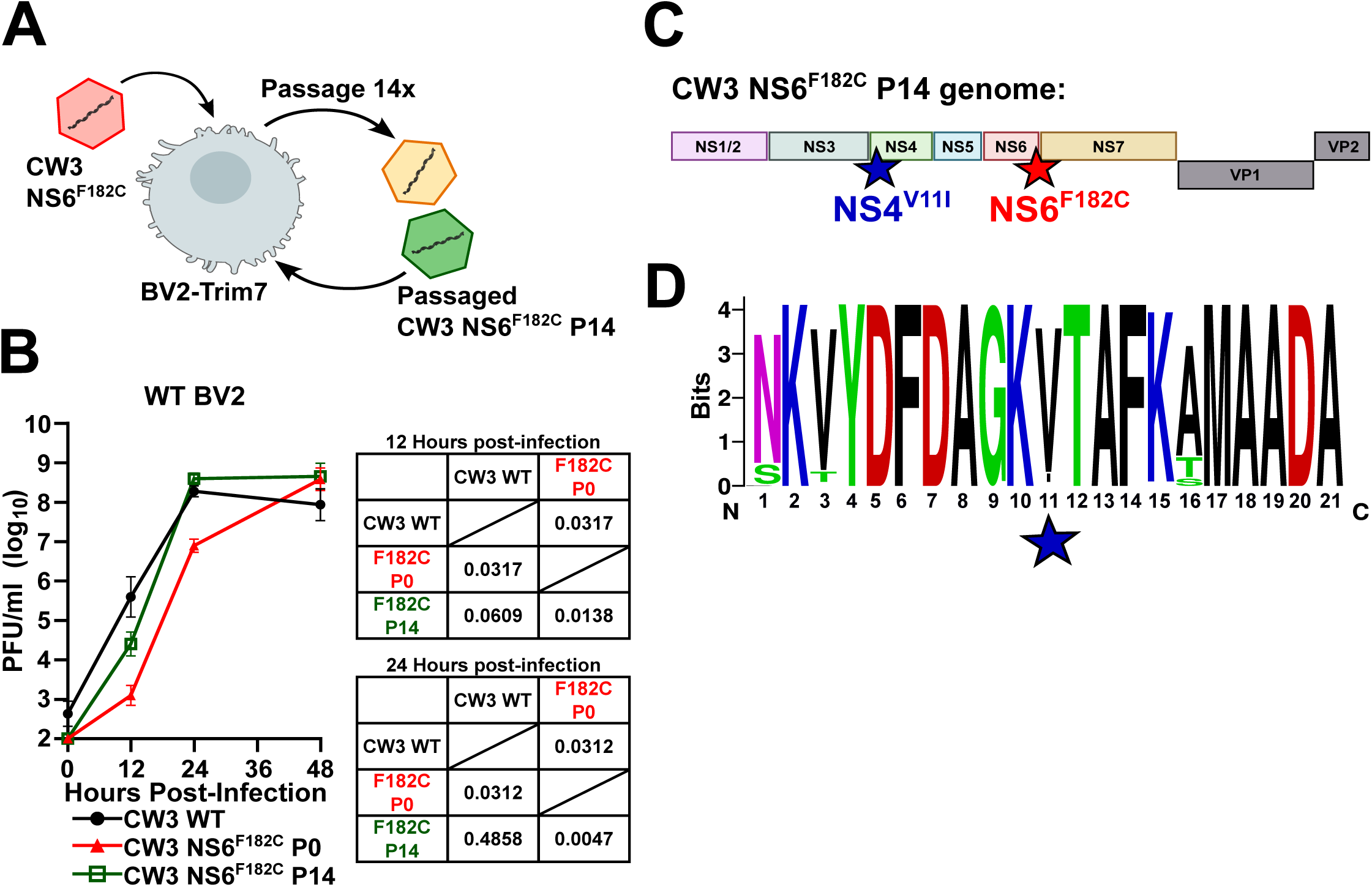
In vitro passaging of MNV^CW3^ NS6^F182C^ rescues its attenuated replication. **A)** Cartoon schematic of the experimental strategy to rescue MNV strain CW3 NS6^F182C^ viral replication. BV2-Trim7 cells were infected with NS6^F182C^ virus and passaged initially at every 48 hours up to passage 6 (P6) and further passaged every 12 hours up to passage 14 (P14). **B)** Wild-type BV2 cells were infected with CW3 wild-type, parental CW3 NS6^F182C^ P0 or passaged CW3 NS6^F182C^ P14 at a multiplicity of infection (MOI) of 0.05. Viral replication was measured using plaque assays at indicated time points. Data is shown as mean ± SEM from three independent experiments. Tables show p-values of statistical comparisons of viral titers compared at 12 hours and 24 hours post-infection. Two-way ANOVA with Tukey’s multiple comparisons test. **C)** Schematic of complete genome of CW3 NS6^F182C^ P14 representing mutations differing from CW3 wild-type. NS6^F182C^ mutation was maintained and an additional mutation in NS4 (NS4^V11I^) was discovered in this passaged virus. **D)** Sequence logo of the first 21 amino acids of NS4 from 87 different MNV strains showing high conservation across all strains. Most MNV strains encode a valine at the 11^th^ position with some strains encoding an isoleucine, similar to the mutation found in the passaged CW3 NS6^F182C^ P14 virus.

To determine the genetic basis for this rescue in replication, we performed consensus sequencing of the P14 viral stock using RT-PCR and sanger sequencing. Importantly, the NS6^F182C^ mutation was maintained in the P14 population, thus we could rule out a genetic reversion for the restored replication (**Figure 1C**). In addition to the F182C, we identified an additional mutation in NS4 leading to a valine to isoleucine substitution (V11I; **Figure 1C**). NS4 is highly conserved in all MNV strains, with most MNV strains encoding a valine-at position 11 while some strains encode an isoleucine (**Figure 1D**). The precise role of NS4 during norovirus replication is unknown as it has no predicted enzymatic activity and no homology to functional domains. Interestingly, two prior investigations identified NS4^V11I^ emerging in viral stocks (12, 13). NS4^V11I^ containing viruses replicated slightly better in vitro than the parental virus, although no other phenotypes were attributed to this mutation (13). Thus, we have uncovered an unappreciated genetic connection between processing of the NS6-7 polyprotein with a mutation in NS4.

### NS4^V11I^ mutation is sufficient to rescue replication of CW3 NS6^F182C^ and wild-type MNV independent of polyprotein processing

To confirm that NS4^V11I^ suppresses the growth defect of NS6^F182C^, we generated a molecular clone containing both variants individually or in combination. A double mutant (NS4^V11I^ + NS6^F182C^) enhanced the overall replication of MNV compared to the NS6^F182C^ mutant alone in BV2 cells (**Figure 2A**). However, this suppression of attenuation only happened at later time points (e.g. 24 hours) as there was no difference in single cycle replication between these two viruses (**Figure 2A**). Interestingly, addition of NS4^V11I^ on a non-attenuated (wild-type) virus had a similar enhancement effect compared to the wild-type virus (**Figure 2A**). These data demonstrate that NS4^V11I^ enhances viral replication of wild-type and NS6^F182C^ containing viruses at later time points of infection.

**Figure 2.**
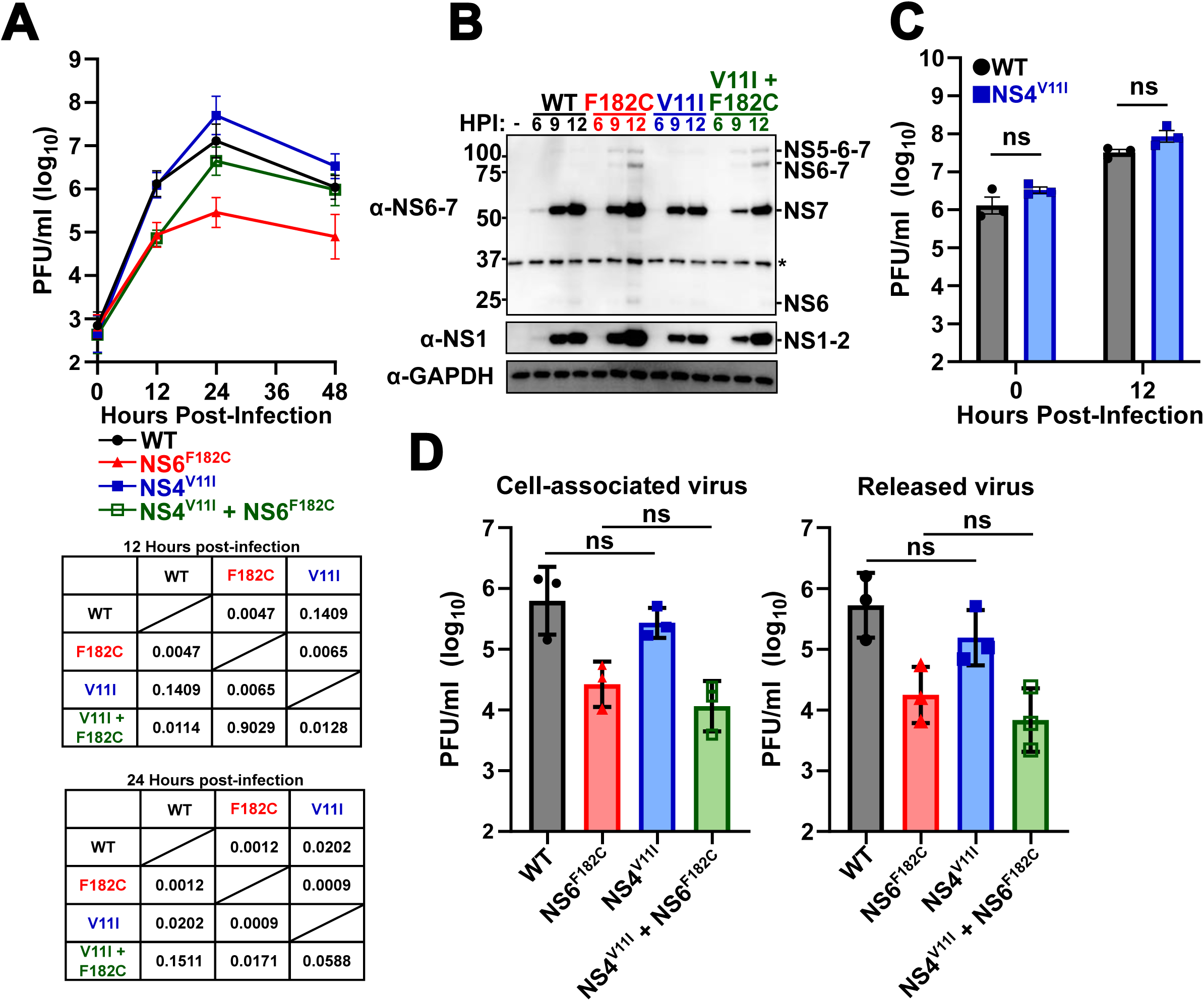
NS4^V11I^ mutation confers rescue of attenuated NS6^F182C^ replication independent of polyprotein processing. **A)** Wild-type BV2 cells were infected with indicated MNV strains derived from MNV molecular clones at a multiplicity of infection (MOI) of 0.05. Viral titers were enumerated using plaque assays at the indicated time points. Data is shown as mean ± SEM from four independent experiments. Tables show p-values of statistical comparisons of viral titers compared at 12 hours and 24 hours post-infection. Two-way ANOVA with Tukey’s multiple comparisons test. **B)** Representative western blot from wild-type BV2 cells infected with indicated MNV strains at an MOI of 5.0 and lysed at the indicated hours post-infection (HPI). NS1/2, NS7 and NS6-7 precursor bands were identified by antibody-specific staining and expected molecular weights of the polyprotein precursors. The asterisk (*) indicates a non-specific band with NS6-7 antibody. **C)** Wild-type BV2 cells were infected with wild-type MNV or MNV NS4^V11I^ at an MOI of 5.0 and viral titers were determined by plaque assay at indicated time points. Data is shown as mean ± SEM from three independent experiments. Two-way ANOVA with Sidak’s multiple comparisons test. **D)** Wild-type BV2 cells were infected with indicated MNV strains at an MOI of 0.05. 12 hours post-infection, cell-associated (representing viral particles within cells) and released virus (representing viral particles in culture supernatants) were harvested. Viral titers were determined by plaque assay. Data is shown as mean ± SEM from three independent experiments. One-way ANOVA with Tukey’s multiple comparisons test.

Position 182 falls within the 3C-protease cleavage site of NS6-7 and mutation of this site to Cysteine (F182C) diminishes, but not abolishes, the cleavage of NS6-7 (9). It is important to note that abolition of the NS6-7 protease cleavage site does not generate infectious MNV (14). Therefore, we tested whether NS4^V11I^ enhances NS6-7 processing which could account for the rescue of viral replication. Consistent with previous findings, viruses containing the F182C mutation had an accumulation in NS6-7 polyprotein precursors including NS6-7 and NS5-6-7 which was not apparent in the wild-type virus (**Figure 2B**). The NS4^V11I^ + NS6^F182C^ double mutant had similar increases in NS6-7 precursor proteins detected during infection (**Figure 2B**). In contrast, the NS6-7 polyprotein pattern of NS4^V11I^ was similar to that of wild-type MNV (**Figure 2B**). Thus, these data indicate that NS4^V11I^ improves viral replication independently of polyprotein processing.

Since we observed an NS4^V11I^-mediated increase in viral replication at 24 hours post-infection but not at 12 hours post-infection (**Figure 2A**), we further tested whether this mutation impacted single-cycle replication. We assessed viral replication at 12 hours post-infection at a high MOI and found no difference in replication between wild-type and NS4^V11I^ (**Figure 2C**). We then assessed whether NS4^V11I^ facilitates faster release of viral particles leading to increase in viral replication over multiple cycles. There was no significant difference in levels of cell-associated virus and released virus at 12 hours post-infection in the NS4^V11I^ viruses relative to their counterparts (**Figure 2D**). Taken together, these data suggest that the NS4^V11I^ improves MNV replication over multiple cycles of replication.

### NS4^V11I^ spreads faster to neighboring cells in culture

We were intrigued by the dramatic increase in viral replication of viruses containing either NS4^V11I^ or the combination of NS6^F182C^ and NS4^V11I^ in multicycle growth curves but not in any measurement of single cycle replication (**Figure 2**). Interestingly, during these experiments, we observed drastic differences in the size of plaques formed by these MNV strains (**Figure 3A**). The specialist, attenuated NS6^F182C^ virus formed very small plaques which were rescued to generalist plaque sizes in the double mutant NS4^V11I^ + NS6^F182C^ virus (**Figure 3A and 3B**). NS4^V11I^ mutant alone also exhibited plaques much larger than wild-type virus (**Figure 3A**). These data suggest an increase in viral spread when NS4^V11I^ mutation is present.

**Figure 3.**
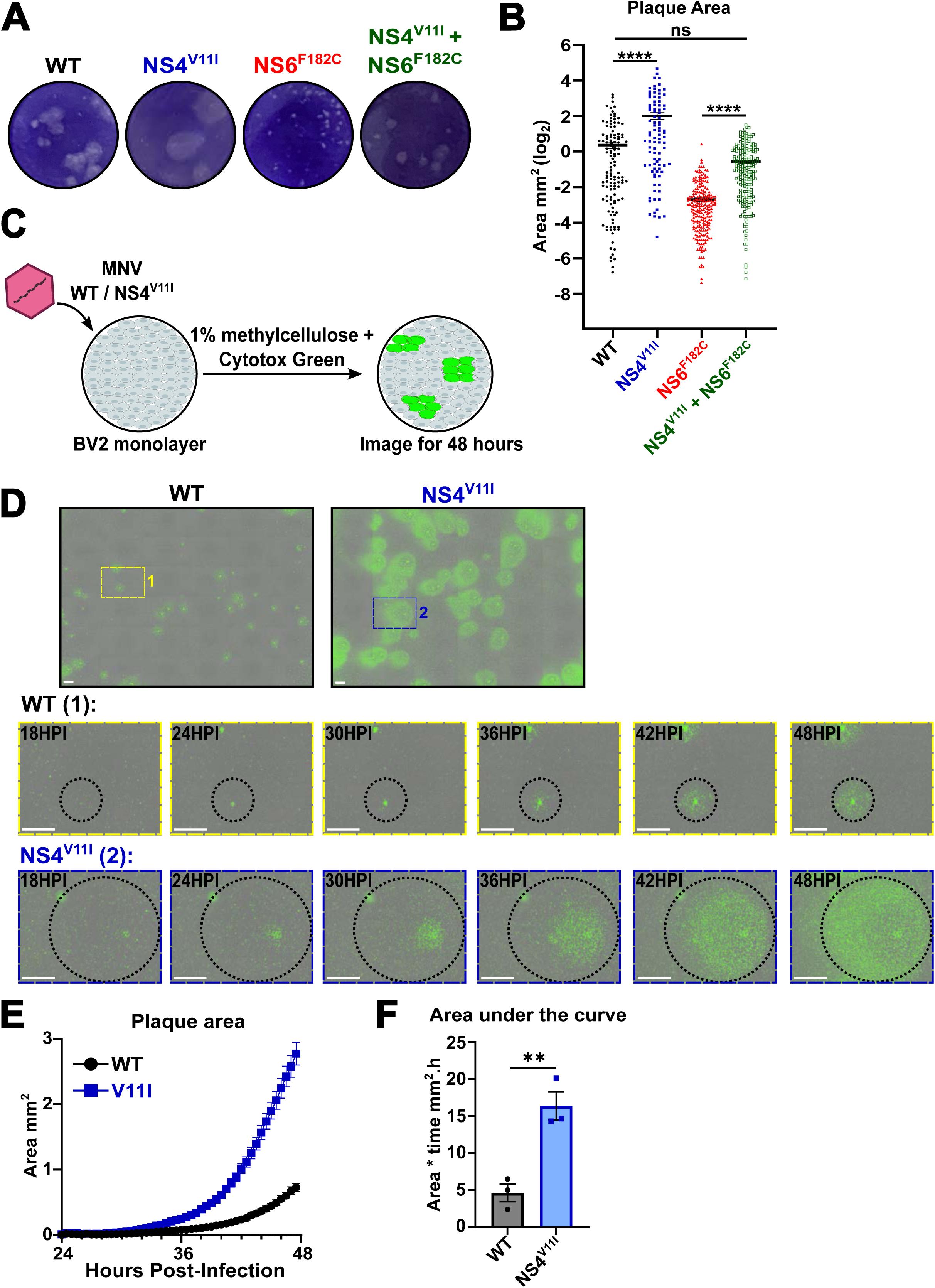
NS4^V11I^ leads to faster viral spread. **A)** Representative plaque assay morphologies of wild-type BV2 cells infected with indicated viral strains. **B)** Quantification of plaque sizes of infections in (A). Plaque areas were measured in ImageJ. Each point represents a single plaque. Data is shown as mean ± SEM from four independent experiments. ns, *P<0.1, **P<0.01, ***P<0.001, ****P<0.0001, Kruskal-Wallis test with Dunn’s multiple comparisons. **C)** Schematic of experimental strategy for real-time imaging of MNV plaque growth. Wild-type BV2 cells were infected with MNV (∼100 pfu per well) and incubated with 1% methylcellulose and 250nM Cytotox green dye. Phase and green-fluorescent images were taken every 2 hours for the first 24 hours post-infection and every 30 minutes for the next 24 hours. **D)** (**Top**) Representative stitched fluorescent microscopy image taken on IncuCyte at 48 hours post-infection of wild-type BV2 cells as infected in (C). Green dye represents apoptotic cells detected by Cytotox Green. (**Bottom**) Time-lapse fluorescence microscopy of selected single plaque growth of wild-type or NS4^V11I^ viral infection over 48 hours-post infection. Scale bar represents 1 mm. **E)** Quantification of plaque area from fluorescent microscopy from the IncuCyte images (D) for 48 hours post-infection. Data represented as mean ± SEM from three independent experiments (n=80-93 plaques per condition). **F)** Area under the curve calculated individually from three independent experiments as in (E). Data represented as mean ± SEM from three independent experiments. **P<0.05, Welch’s t-test.

To quantify the kinetics of viral spread and plaque formation, we designed an assay for real-time measurement of MNV plaque formation (**Figure 3C**). This assay incorporates a very low infection (100 plaque forming units per well) with the viscous, liquid overlay methylcellulose to limit viral diffusion. Additionally, we supplemented the media with Cytotox Green dye which marks dead cells and can be used as a surrogate for viral replication as there is no real-time live monitoring system for MNV replication. We imaged the cells over 48 hours. We focused our efforts on characterizing the differences between wild-type MNV and MNV containing NS4^V11I^. These two viruses both produce robust, easily detectable plaques and demonstrate an NS4^V11I^-mediated enhancement of viral spread. Using Cytotox Green to mark plaques, we observe nearly a 3-fold difference in plaque size after 48 hours, consistent with our crystal violet-based measurements (**Figure 3B, 3D, and 3E**). Plaque initiation was detectable at 24 hours post-infection (HPI) for both wild-type and NS4^V11^ containing viruses (**Figure 3E**). Despite the initiation event occurring at a similar time point, we observed a significant increase in spread of the NS4^V11I^ viral plaques over time (**Figure 3D and 3E**). Quantification of nearly 100 plaques in each condition across multiple experiments demonstrated that despite heterogeneity in plaque area within a sample, the NS4^V11I^ plaques were larger and spread faster (**Figure 3E**). The area under the curves from three independent experiments was significantly greater in NS4^V11I^ compared to wild-type infection demonstrating reproducibility across independent experiments (**Figure 3F**). Taken together, these data demonstrate that the NS4^V11I^ containing viruses are able to spread to neighboring cells significantly faster and likely accounts for the suppression of the NS6^F182C^ attenuating phenotype.

### Rapid CW3 NS4^V11I^ viral spread is not due to modulation of interferon signaling

Since NS4^V11I^ mutation enables a more rapid spread of infection from a single infected cell to its neighboring cells, we hypothesized that NS4^V11I^ may be modifying host defense pathways to infect neighboring cells more easily. Type I and type III interferons (IFN) are a major innate immune defense against viruses, including norovirus (15–18). To determine whether the NS4^V11I^ mutation improves viral spread to neighboring cells due to modulation of interferon signaling, we compared the plaque size of viruses grown on wild-type BV2 or BV2ΔSTAT1, which lack the ability to respond to all IFNs. There was no difference in the plaque morphology or size of wild-type MNV in wild-type or BV2ΔSTAT1 cells (**Figure 4A and 4B**). We observed similar trends in viruses containing the NS4^V11I^ mutation (**Figure 4A and 4B**). Interestingly, viruses that contained the NS6^F182C^ mutation had an increase in plaque size when grown on BV2ΔSTAT1 cells compared to wild-type BV2 cells (**Figure 4A and 4B**). Importantly, the NS4^V11I^ increase in viral spread as measured by plaque size occurred independently of STAT1 dependent signaling (**Figure 4A and 4B**). For example, the NS4^V11I^ and NS6^F182C^ double mutant had an increase in plaque size compared to NS6^F182C^ single mutant in both wild-type and BV2ΔSTAT1 cells (**Figure 4A and 4B**). These data demonstrate that the increase in viral spread by the NS4^V11I^ does not occur via modulation of the IFN pathway but likely through an unappreciated host-defense pathway.

**Figure 4.**
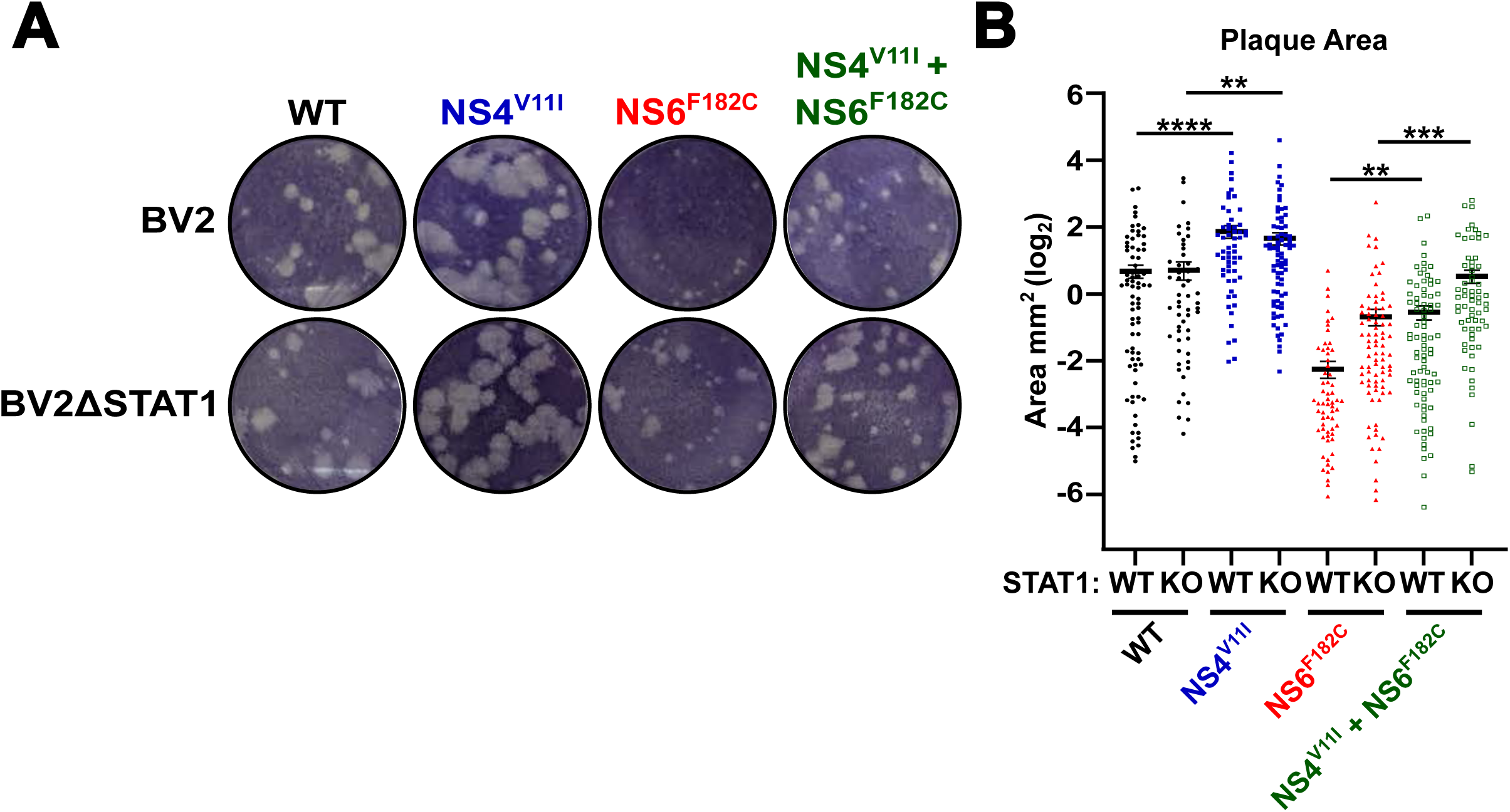
Rapid spread of MNV NS4^V11I^ is not due to modulation of interferon signaling. **A)** Representative plaque assay morphologies of wild-type BV2 (top) and BV2ΔSTAT1 (bottom) infected with indicated MNV strains. **B)** Quantification of plaque sizes of infections in (A). Plaque areas were measured in ImageJ. Each point represents a single plaque. Data is shown as mean ± SEM from four independent experiments. ns, *P<0.1, **P<0.01, ***P<0.001, ****P<0.0001, Kruskal-Wallis test with Dunn’s multiple comparisons.

## Discussion

Viruses are a powerful genetic tool to study evolution due to their compact genome and prolific replication within a relatively short time frame. These traits enable viruses to rapidly adapt to new environments. Studying these adaptations has enabled the development of live attenuated vaccines, model zoonotic transmission, discovery of immune barriers, and identification of antiviral targets (1, 2, 19). In many cases these adaptations lead to specialization in a particular environment and a loss of general replication in other settings. While genetic reversion can allow these specialized viruses to become generalists, there is less information about how suppressive mutations enable a specialist virus to return to the original generalist state. Here, using an unbiased forward genetic screen, we describe such a situation in which a suppressive mutation is acquired to restore the replication of a specialist MNV strain to the levels of its original precursor. MNV adapted to Trim7 overexpressing cells (MNV NS6^F182C^) can no longer grow robustly in wild-type or other general settings (9). However, the acquisition of a mutation in NS4 (Valine to Isoleucine at position 11; V11I) converts the attenuated, specialist virus to a robustly replicating generalist (**Figure 1 and 2**). These findings have several implications on norovirus biology and viral evolution.

Viruses within the Caliciviridae family differ in their polyprotein processing of NS6 and NS7 (also known as Pro-Pol) (20, 21). Norovirus is unique amongst caliciviruses for its separation of NS6 and NS7, and surprisingly this cleavage event is required for recovery of infectious virus (14). In support of this notion, the NS6^F182C^ mutation, which has decreased cleavage of the NS6-7 precursor, is positively selected as a means of resisting Trim7 expression yet is highly attenuated in native settings (9). It remains unclear why the cleavage of NS6-7 is uniquely required in noroviruses. Our data suggests one mechanism may be that cleaved, norovirus NS6 is more efficient at antagonizing innate immunity and is necessary for efficient viral spread. In support of this model, we find that a single amino acid mutation in MNV NS4 (NS4^V11I^) is sufficient to rescue the attenuated replication of NS6^F182C^. This mutation did not alter the defective polyprotein processing of NS6^F182C^ but rather increases the spread of both the parental virus and the F182C attenuated virus (**Figure 2**). Increased viral spread via NS4^V11I^ may compensate for a decrease in NS6-mediated innate immune antagonism. In support of this hypothesis, NS6 has several reported immune antagonism functions including alteration of host translation through cleavage of polyA-binding protein (PABP) (22), and cleavage of NF-κB essential modulator (NEMO) protein (23). Additionally, other critical immune nodes are targeted by viral 3C-proteases, including by related caliciviruses (24). However, it is also possible that NS6^F182C^ has multiple bottlenecks in viral replication and the NS4^V11I^ is a simple method to significantly boost viral spread and increase fitness.

The function of NS4 during norovirus replication is still unknown. Based on positional homology in the genome, NS4 is a predicted “3A-like” protein similar to picornavirus 3A protein. 3A proteins have diverse functions but can be largely classified as membrane remodelers and immune antagonists. For example, poliovirus 3A regulates membrane remodeling in host cells (25) and inhibits cellular protein secretory pathways to decrease inflammatory immune response (26, 27). Indeed, ectopic expression studies of human and mouse norovirus NS4 proteins have demonstrated a propensity for membrane rearrangement (28), possible alterations in the general secretory pathway (29), and the potential for the activation of innate immunity through sensing cytosolic DNA (30). However, the physiological role of NS4 during infection remains enigmatic. It is possible that the comparison of the Valine to Isoleucine switch in NS4 during infection may provide novel insights into the mechanism by which norovirus utilizes NS4. Future studies may leverage the significant difference in plaque sizes of NS4 variants as a robust platform to dissect the host and viral pathways enabling increased viral spread.

Position 11 of NS4 amongst MNV strains is predominantly valine, although a significant number of strains have an isoleucine at this position. Our in vitro data demonstrates that NS4^V11I^ replicates to higher titers and spreads better in vitro, yet a previous study found no difference in pathogenesis in STAT1^-/-^ mice infected with either variant (13). It remains unclear what evolutionary pressures differ in vitro and in vivo to have such differences in selection and maintenance of these different variants. Valine and isoleucine are nearly identical chemically, yet the viral spread of NS4^V11I^ is dramatic and robust (**Figure 3)**. Future work investigating the molecular basis for these different phenotypes will likely increase our understanding of norovirus evolution and the function of NS4.

## Materials and Methods

### Cell culture

BV2, BV2-ΔSTAT1 (Dr. Skip Virgin, Washington University; (31)) and HEK-293T cells (ATCC) were cultured in Dulbecco’s Modified Eagle Medium (DMEM) supplemented with 5% fetal bovine serum (FBS). Trim7-expressing stable cell lines were generated by lentiviral transduction. Briefly pCDH-MCS-T2A-Puro-MSCV-Trim7 isoform 1 (8), lentiviral packing vector (psPax2) and psuedotyping vector (pCMV-VSV-G) were co-transfected into HEK-293T cells using OptiMEM (Sigma) and Transit-LT1 (Mirus). Lentivirus was collected 48 hours post-transfection, filtered through a 0.45um filter and added to cells. 48 hours post-transduction, cells were selected with 1ug/mL puromycin.

### MNV assays

MNV^CW3^ stocks were generated from plasmids encoding the complete MNV^CW3^ genome (GenBank ID EF014462.1), purified and titered by plaque assay as described previously (32, 33). NS4^V11I^ and NS6^F182C^ mutations were introduced through splicing by overlap extension PCR in MNV molecular clones. All plasmid sequences were verified through sequencing prior to use. Mutant viruses were generated on the plasmid containing the MNV^CW3^ backbone similar to parental MNV^CW3^ stocks. Genetic identity of viral stocks was confirmed by sequencing viral RNA. MNV replication assays were performed as described previously (8) with the indicated multiplicity of infection (MOI 0.05 or MOI 5). MNV protease processing assay was performed by infecting WT BV2 cells at an MOI of 5. Cells were lysed at indicated times post-infection in Laemmli buffer (Bio-Rad). Lysates were resolved via SDS-PAGE, transferred to polyvinylidene difluoride (PVDF) membranes and probed with the indicated antibodies.

To determine the cell-associated and released virus quantities, WT BV2 cells were seeded at 2.5 × 10^5^ per well of a 24-well plate. Cells were allowed to adhere overnight and infected with MNV^CW3^ WT and mutant viruses at an MOI of 0.05. 12 hours post-infection, supernatants were collected from the wells, and fresh media was added back to the wells. Both samples were frozen at -80°C. Viral titers were determined by plaque assay.

Viral spread plaque assays were performed by seeding WT BV2 or BV2ΔSTAT1 at 2.5 × 10^5^ cells per well in a 24-well plate. After adhering overnight, cells were infected with a serial dilution of 10, 100, and 1000 PFU of the indicated viruses. After 1 hour of incubation, virus was aspirated and 1% methylcellulose was added to the wells. Cells were stained with crystal violet at 48 hours post-infection and stained plates were imaged. Quantification of plaque sizes was performed in ImageJ.

### Viral evolution

MNV passaging experiments were conducted in BV2-Trim7 cells similar to our previous study (9). Briefly, BV2-Trim7 cells were seeded at 1 × 10^7^ in a 10cm^2^ plate. After adhering overnight, attenuated parental CW3 NS6^F182C^ (P0) was infected at MOI 5. 24 hours post-infection, virus was harvested and clarified from the supernatant to obtain CW3 NS6^F182C^ P1. 1mL of the P1 virus was added to BV2-Trim7 cells in a 10cm^2^ plate. Subsequent passaging of the virus was done at 48 hours post-infection up to passage 6 (P6). After passage 6, viral passaging was continued at every 12 hours post-infection up to passage 14 (P14). This provided more evolutionary selective pressure to the virus. 1mL of the final clarified P14 viral supernatant was used to isolate total RNA using Zymo Direct-zol kit. cDNA was synthesized using the M-MLV Reverse Transcriptase kit (Invitrogen) following manufacturer’s protocols. Fragment PCR was performed on the viral cDNA using primers spanning the MNV genome. The PCR products were then purified and sequenced by Sanger sequencing.

### Antibodies and Western blotting

Rabbit polyclonal anti-NS6-7 (a kind gift from Kim Green), mouse monoclonal anti-NS1 (a kind gift from Sanghyun Lee), mouse anti-GAPDH HRP (Sigma-Aldrich), anti-Rabbit IgG HRP (Sigma-Aldrich), and anti-mouse IgG (Sigma-Aldrich) were used for western blotting.

### IncuCyte-based real time quantification of MNV plaques

WT BV2 cells were seeded at 2 × 10^6^ per well in a 6-well plate. After adhering overnight, cells were infected with CW3 WT or CW3 NS4^V11I^ at 100 PFU per well. After one hour of incubation, virus was aspirated from the wells. 1% methylcellulose supplemented with 250nM Cytotox Green dye (Sartorius) was added to the wells. The methylcellulose was first spun down at 4000 rpm for 10 min to remove any particulates that may interfere with imaging. The plate was placed in the IncuCyte and images were acquired in the phase and green channels. 36 images were taken per well at 4x magnification at every 2 hours up to 24 hours post-infection and every 30 minutes for the next 24 hours. Timelapse videos of each image section was exported and stitched together using Adobe PremierePro. Videos were processed on ImageJ and plaque area of selected plaques were calculated over time for each experiment.

## Acknowledgements

We would like to thank Dr. Aparijita Dasgupta and all members of the Orchard lab for helpful discussions on this project. This work was supported by a grant from the NIH to R.C.O. (5R01DK133231).

## Author Contributions

M.A. S. designed the project, performed experiments, analyzed data, and wrote the paper. R.C.O. conceptualized the project, provided supervision, and wrote the paper. All authors read and edited the manuscript.

